# Disrupted copper availability in sporadic ALS: Implications for Cu^II^(atsm) as a treatment option

**DOI:** 10.1101/2020.04.17.047704

**Authors:** James BW Hilton, Kai Kysenius, Jeffrey R Liddell, Carsten Rautengarten, Stephen W. Mercer, Bence Paul, Joseph S Beckman, Catriona A. McLean, Anthony R White, Paul S Donnelly, Ashley I Bush, Dominic J Hare, Blaine R Roberts, Peter J Crouch

## Abstract

**Objective:** The copper compound Cu^II^(atsm) is in phase 2/3 testing for treatment of the neurodegenerative disease amyotrophic lateral sclerosis (ALS). Cu^II^(atsm) consistently and reproducibly ameliorates neurodegeneration in mutant SOD1 mouse models of ALS where its neuroprotective activity has been ascribed in part to improving availability of copper to essential cuproenzymes. However, SOD1 mutations cause only ~2% of ALS cases with most cases being of unknown aetiology. Therapeutic pertinence of Cu^II^(atsm) to sporadic ALS is therefore unclear.

**Methods:** We assayed post-mortem spinal cord tissue from sporadic cases of ALS for the anatomical and biochemical distribution of copper, the expression of genes involved in copper handling, and the activities of cuproenzymes.

**Results:** The natural distribution of copper is disrupted in sporadic ALS. The ALS-affected tissue has a molecular signature consistent with an unsatiated requirement for copper and cuproenzyme activity is affected. Copper levels are decreased in the ventral grey matter, the primary anatomical site of neuronal loss in ALS.

**Interpretation:** Mice expressing mutant SOD1 recapitulate salient features of ALS. The unsatiated requirement for copper that is evident in these mice is a biochemical target for Cu^II^(atsm). Evidences provided here for disrupted copper bioavailability in human cases of sporadic ALS indicate that a therapeutic mechanism for Cu^II^(atsm) involving copper bioavailability is pertinent to sporadic cases of ALS, and not just those involving mutant SOD1.

## INTRODUCTION

The orally bioavailable and blood-brain barrier penetrant copper compound diacetyl-bis(4-methylthiosemicarbazonato)copper^II^ [Cu^II^(atsm)] is one of the most robustly tested and corroborated candidate drugs ever developed for the neurodegenerative condition of ALS, a fatal and rapidly progressive disease that selectively destroys motor neurones in the central nervous system (CNS). Pre-clinical studies involving mouse models of the disease show that oral treatment with Cu^II^(atsm) protects motor neurones in the spinal cord, mitigates the motor symptoms of neuronal decline, and extends survival^1–7^. These outcomes include mitigation of disease progression when the treatment commences after symptom onset^2^. In animal model testing, Cu^II^(atsm) outperformed riluzole, the current first-line clinical prescription^2^. Cu^II^(atsm) is the first and only candidate drug for ALS to be independently validated by the ALS Therapy Development Institute^6^.

Copper is an essential micronutrient and its redox cycling between oxidised Cu^II^ and reduced Cu^I^ mediates catalytic activity in numerous vital cuproenzymes. The first discovered genetic cause of ALS was autosomal-dominant mutation of superoxide dismutase 1 (SOD1)^8^, a ubiquitously expressed cuproenzyme antioxidant. Transgenic mice expressing mutant SOD1 are a robust animal model of ALS, possessing face and construct validity^9^. Overexpression of SOD1 in these models generates a pool of SOD1 that lacks copper in the catalytic site^1,3,5,10^. Treatment with Cu^II^(atsm) enables copper from the complex to become incorporated into SOD1^3^, thereby converting aberrant copper-deficient SOD1 to its physiologically mature copper-replete form.

These outcomes support a protective mechanism for Cu^II^(atsm) implicating its ability to safely increase copper bioavailability in the CNS. Insufficient copper bioavailability in these animals is a CNS-specific defect despite ubiquitous expression of the disease-causing gene^10^. The copper-dependent activities of cuproenzymes other than SOD1 are also affected^5,11^, indicating the causal gene has a broad impact on copper homeostasis. However, SOD1 mutations account for only ~2% of ALS cases in the clinic, with most cases being sporadic and of unknown aetiology. Whether the efficacy of Cu^II^(atsm) in mutant SOD1 mice could translate to sporadic ALS through a mechanism of action involving copper bioavailability, remains unclear. To this end, here, we assessed human spinal cord tissue from sporadic cases of ALS for levels of copper, cuproenzyme activities and the expression of regulatory pathways.

## SUBJECTS/MATERIALS AND METHODS

### Human tissue and processing

Procedures involving post-mortem human tissue (Table 1) were approved by a University of Melbourne Human Ethics Committee (Project ID 1238124). Frozen sections of lumbar spinal cord were obtained from the Victorian Brain Bank (Australia) and the MS Society Tissue Bank (UK). Tissue used in microdroplet and biochemical analyses was processed to generate tris(hydroxymethyl)aminomethane-buffered saline (TBS)-soluble and -insoluble fractions as previously described^11^. Assays involving ceruloplasmin, dopamine β-hydroxylase (DβH) and lysyl oxidase utilised TBS-insoluble fractions that were supplemented with 1% (v/v) Triton X-100 then centrifuged (18,000 RCF, 4°C, 5 min) to produce Triton X-100 soluble extracts. Grey matter material dissected from spinal cord samples was processed for gene expression as described below.

**Table 1.** Sex, age and post-mortem interval details for all study control and ALS cases.

| Sample code | Disease group | Sex | Age | PMI (hrs) |
| --- | --- | --- | --- | --- |
| C1 | Control | Male | 52 | 33 |
| C2 | Control | Male | 64 | 68 |
| C3 | Control | Male | 86 | 71 |
| C4 | Control | Female | 89 | 6 |
| C5 | Control | Male | 76 | 50 |
| C6 | Control | Male | 85 | 68 |
| C7 | Control | Female | 78 | 23 |
| C8 | Control | Female | 89 | 22 |
| C9 | Control | Male | 68 | 30 |
| C10 | Control | Male | 90 | 12 |
| C11 | Control | Male | 66 | 54 |
| C12 | Control | Female | 89 | 13 |
| ALS1 | Sporadic ALS | Female | 56 | 45 |
| ALS2 | Sporadic ALS | Female | 80 | 23 |
| ALS3 | Sporadic ALS | Male | 60 | 16 |
| ALS4 | Sporadic ALS | Male | 73 | 21 |
| ALS5 | Sporadic ALS | Male | 65 | 45 |
| ALS6 | Sporadic ALS | Male | 72 | 67 |
| ALS7 | Sporadic ALS | Male | 75 | 40 |
| ALS8 | Sporadic ALS | Male | 79 | 31 |
| ALS9 | Sporadic ALS | Male | 64 | 14 |
| ALS10 | Sporadic ALS | Female | 77 | 25 |
| ALS11 | Sporadic ALS | Female | 65 | 16 |
| ALS12 | ALS* | Male | 80 | 25 |
| ALS13 | Sporadic ALS | Male | 83 | 36 |
| ALS14 | Sporadic ALS | Male | 58 | 22 |
| ALS15 | Sporadic ALS | Male | 55 | 56 |
| ALS16 | Sporadic ALS | Female | 59 | 20 |
| ALS17 | Sporadic ALS | Male | 54 | 55 |
| ALS18 | Sporadic ALS | Female | 55 | 28 |
| ALS19 | Sporadic ALS | Female | 59 | 58 |
| ALS20 | Sporadic ALS | Male | 71 | 30 |
| ALS21 | Sporadic ALS | Male | 69 | 36 |
| ALS22 | Sporadic ALS | Male | 65 | 22 |
| ALS23 | Sporadic ALS | Male | 71 | 22 |
| ALS24 | Sporadic ALS | Male | 68 | 66 |

\*Adopted, no known genetic history of ALS. PMI: Post-mortem interval.

### Mouse tissue

Procedures involving mice were approved by a University of Melbourne Animal Experimentation Ethics Committee (Project ID 1111947) and conducted in accordance with National Health and Medical Research Council guidelines. Spinal cords were collected from the SOD1^G37R^ model of ALS^12^ (JAX Stock # 008342) and non-transgenic littermate controls as previously described^11^. Mice were 180-190 days old, which equates to mid-to late-stage symptom development^11^. Lumbar sections of cord were processed for gene expression as described below.

### Copper quantitation

*In situ* quantitation of copper was performed using laser ablation-inductively coupled plasma-mass spectrometry (LA-ICP-MS)^13^ utilising spinal cord embedded in Optimal Cutting Temperature compound and cryo-sectioned at 30 µm in the transverse plane. Total copper levels in human spinal cord were measured by solution ICP-MS as previously described^1^ using an Agilent 7700 Series ICP-MS. To assess biochemical partitioning of copper, TBS-soluble and TBS-insoluble fractions were analysed using microdroplet LA-ICP-MS^14^. All *in situ* and microdroplet LA-ICP-MS analyses utilised a NewWave Research NWR213 laser ablation system coupled to an Agilent 8800 triple quadrupole ICP-MS. Data were analysed using Iolite operating under the Igor Pro 8 suite (WaveMetrics, Inc.).

### Gene expression

Samples were prepared using manufacturer’s instructions as follows: Nucleic acids were extracted (TRI-Reagent, Sigma); contaminating gDNA in isolated mRNA was degraded (Turbo DNA-Free Kit, Thermo Fisher Scientific); mRNA quantity was measured (Qubit RNA HS Assay Kit, Thermo Fisher Scientific); cDNA was synthesised (High Capacity cDNA Reverse Transcription Kit, Thermo Fisher Scientific); and cDNA (25ng) was pre-amplified for all genes assessed (Taqman PreAmp Master Mix and Taqman Gene Expression Assays, Thermo Fisher Scientific). Pre-amplified cDNA was diluted 20-fold, then quantitative RT-PCR performed (Taqman Gene Expression Assays and Taqman Fast Advanced Mastermix, Thermo Fisher Scientific) on samples in triplicate using a QuantStudio 6 Flex system (Thermo Fisher Scientific). Relative expression of individual genes involved in cellular copper handling was determined via the ΔΔct method normalised to *GAPDH/Gapdh*. Composite z-score values representing changes affecting overall copper handling were calculated for each control or ALS case (or animal) as the average z-score across all genes analysed.

### SDS-PAGE and immunoblotting

Proteins were resolved by SDS-PAGE and assessed by immunoblotting using methods previously described^11^. Primary antibodies used were raised to detect: SOD1 (Abcam, ab79390); GAPDH (Cell Signaling Technology, 2118); ceruloplasmin (DAKO, Q0121); DβH (NBP1-31386, Novus); lysyl oxidase (Abcam, 174316); and lysyl oxidase homologue 3 (ARP60280-P050, Aviva Systems Biology). Detection utilised horseradish peroxidase conjugated secondary antibodies for anti-rabbit IgG (Cell Signaling Technology, 7074) or anti-mouse IgG (Cell Signaling Technology, 7076) followed by enhanced chemiluminescence (ECL Advance, GE Healthcare). Abundance of cuproproteins of interest was normalised to the loading control GAPDH and expressed relative to control cases.

### Cuproenzyme activity

SOD1 activity in the TBS-soluble fraction and ceruloplasmin ferroxidase activity in Triton X-100 extracts were determined as previously described^11^.

DβH activity was measured using liquid chromatography tandem mass spectrometry (LC-MS/MS) to monitor enzymatic production of the DβH product norepinephrine using a procedure based on a previous report^15^. Triton X-100 extracts were combined with a reaction mixture containing 200 mM sodium acetate (pH 5.0), 30 mM N-ethylmaleimide, 5 µM CuSO_4_ (or equivalent volume of dH_2_O), 870 U catalase, 10 mM sodium fumarate and 10 mM ascorbate. Following pre-incubation (37°C, 5 min), reactions were initiated by adding 10 mM dopamine and then incubated at 37°C for 45 minutes.

Epinephrine was added to each sample as an internal standard, followed by 2 mL 100 mM NH_4_H_2_PO_4_ (pH 10.0) supplemented with 2% (v/v) stabiliser (500 mM EDTA, 317 mg mL^-1^ sodium metabisulfite). Samples were then subjected to solid phase extraction using Bond Elut phenylboronic acid (PBA) 100 mg, 3 mL cartridges (Agilent). Cartridges were equilibrated with 1 mL acetonitrile then 1 mL 5% (v/v) formic acid in methanol, then 1 mL of 100 mM NH_4_H_2_PO_4_ (pH 10.0) supplemented with 2% (v/v) stabiliser. After sample addition, another 1 mL 100 mM NH_4_H_2_PO_4_ (pH 10.0) supplemented with 2% (v/v) stabiliser was added. The matrix was washed sequentially with 2 mL 1% (v/v) NH4OH in 95% (v/v) methanol, 2 mL of 1% (v/v) NH4OH in 95% (v/v) acetonitrile, then 1% (v/v) NH4OH in 30% (v/v) acetonitrile. Samples were dried under vacuum then analytes eluted using 3 × 500 µL aliquots of 5% (v/v) formic acid in methanol then evaporated in a vacuum concentrator before reconstitution in 0.3% (v/v) formic acid in dH_2_O.

LC-MS/MS analyses were performed using an Agilent 1100 series HPLC system and a Sciex 4000 QTRAP LC-MS/MS system equipped with a TurboIonSpray ion source. The system was run in Micro mode using a mix rate of 400 µL min^-1^ with the column compartment set to 50°C and samples kept at 20°C. Catecholamine analytes were separated using a Hypercarb column (150 × 1 mm, 5 µm particle size, Thermo Fisher Scientific) at a flow rate of 50 µL min^-1^. Initial run conditions used 99% buffer A (0.3% (v/v) formic acid in dH_2_O) and 1% buffer B (100% acetonitrile) for 1 minute followed by a gradient to 25% buffer B within 20 minutes, then 80% buffer B within 2 minutes. Conditions were then held at 80% buffer B for 2 minutes before a return to 1% buffer B within 2 minutes and holding at 1% buffer B for 6 minutes.

The QTRAP was set to positive ion mode using the multiple reaction monitoring (MRM) scan type. Conditions were: spray voltage 4200V; source temperature 425°C; collision gas set to high; source gas 1 and source gas 2 set to 20. A time of 100 ms was applied to each transition resulting in a duty cycle of 1.0501 seconds, with Q1 and Q3 resolutions set to Unit. Data were collected using the Analyst 1.5.1 Build 5218 (Sciex) operating in MRM mode. Catecholamine analytes were quantified using the MultiQuant 2.1 (build 2.1.1296.02.1) software package (Sciex) through integration of signal peaks for norepinephrine, dopamine and epinephrine. DβH activity was calculated as amount of norepinephrine produced min^-1^ mg^-1^ sample protein.

Lysyl oxidase activity was measured via production of fluorescent resorufin from the substrate Amplex UltraRed (Thermo Fisher Scientific) based on a previously established protocol^16^. Assay buffer (50 mM sodium borate, 1 M urea, 10 mM CaCl_2_, pH 8.0) was used to make a 4 mM benzylamine (Sigma) solution. Equal volumes of Triton X-100 soluble sample in assay buffer and benzylamine solution were then loaded into a 96-well plate. After incubation (37°C, 30 min), each well was loaded with an equivalent volume of substrate mixture (2 U mL^-1^ horseradish peroxidase and 40 µM Amplex UltraRed in assay buffer) then fluorescence measured (Ex. 544 nm, Em. 590 nm). Lysyl oxidase activity was calculated relative to recombinant lysyl oxidase protein standards (OriGene).

### Statistical analyses

All statistical analyses were performed using GraphPad Prism. Data sets were assessed for statistical outliers using Iterative Grubbs’ (α≤0.05). Data are presented as mean ±S.E.M. except for transcript data which are presented as z-scores. Significant differences between groups were determined using two-tailed t-tests except for data presented in Fig. 5C which involved two-tailed paired t-tests. Significance was determined as P<0.05.

## RESULTS

### Copper distribution is altered in the sporadic ALS affected spinal cord

Total copper levels were unaltered in whole homogenates of the ALS affected spinal cord (Fig. 1A). However, analysis of TBS-soluble and TBS-insoluble fractions via our microdroplet method^14^ revealed partitioning of endogenous copper in sporadic ALS was shifted towards the insoluble fraction (Fig. 1B-D). In addition, our *in situ* quantitation of copper by LA-ICP-MS^13,17,18^ revealed that in ALS endogenous copper was decreased in the grey matter and increased in the white matter, with the ventral horn grey matter and dorsolateral white matter regions being the principal anatomical sites of decrease and accumulation respectively (Fig. 2, Table 2). The ventral horn grey matter is the region within the spinal cord where the soma of motor neurones primarily reside and the corticospinal tracts traverse through the dorsolateral white matter column. Thus, these primary anatomical sites of pathology in ALS display the most conspicuous changes in copper.

**Table 2.**
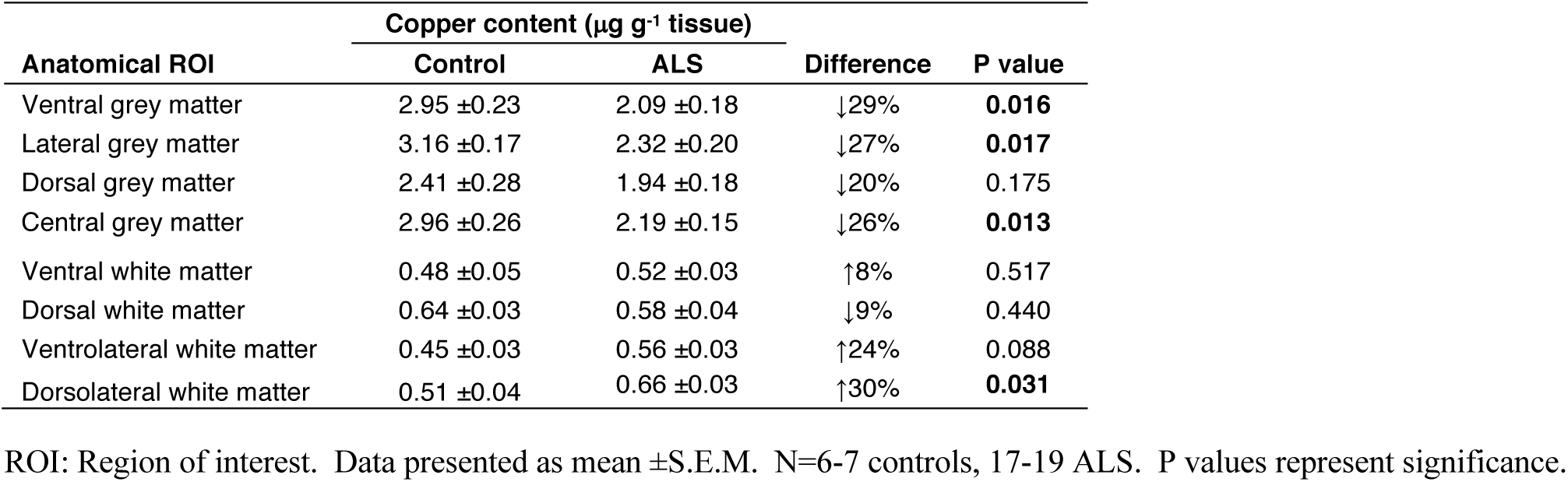
Copper content in anatomical sub-regions of human spinal cord.

| **Anatomical ROI** | **Copper content ($\mu\text{g g}^{-1}$ tissue)** |  | **Difference** | **P value** |
| --- | --- | --- | --- | --- |
|  | **Control** | **ALS** |  |  |
| Ventral grey matter | 2.95 ±0.23 | 2.09 ±0.18 | ↓29% | **0.016** |
| Lateral grey matter | 3.16 ±0.17 | 2.32 ±0.20 | ↓27% | **0.017** |
| Dorsal grey matter | 2.41 ±0.28 | 1.94 ±0.18 | ↓20% | 0.175 |
| Central grey matter | 2.96 ±0.26 | 2.19 ±0.15 | ↓26% | **0.013** |
| Ventral white matter | 0.48 ±0.05 | 0.52 ±0.03 | ↑8% | 0.517 |
| Dorsal white matter | 0.64 ±0.03 | 0.58 ±0.04 | ↓9% | 0.440 |
| Ventrolateral white matter | 0.45 ±0.03 | 0.56 ±0.03 | ↑24% | 0.088 |
| Dorsolateral white matter | 0.51 ±0.04 | 0.66 ±0.03 | ↑30% | **0.031** |

ROI: Region of interest. Data presented as mean ± S.E.M. N=6-7 controls, 17-19 ALS. P values represent significance.

**Figure 1.**
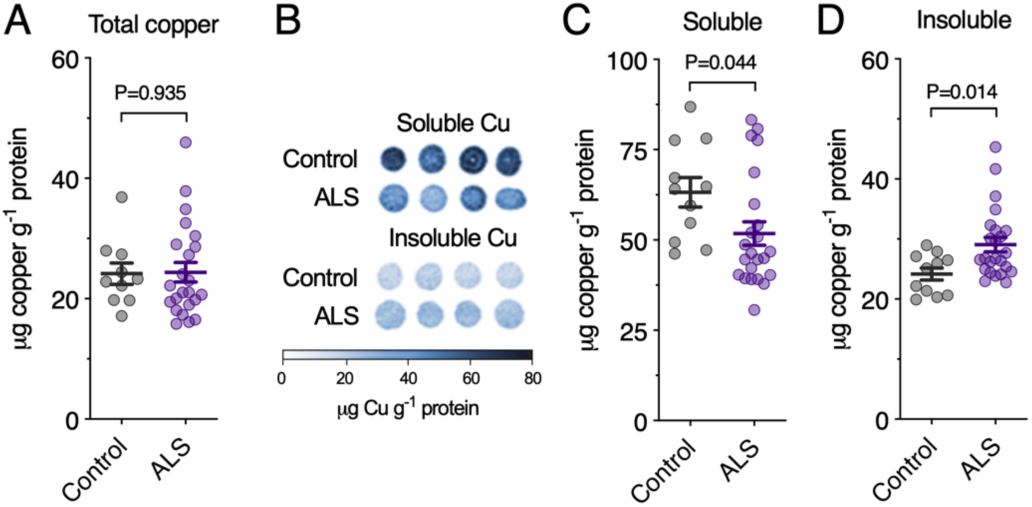
Copper in sporadic ALS-affected spinal cord relative to biochemical partitioning. **A**, Copper in whole homogenate of human spinal cord prepared in tris-buffered saline (TBS) based extraction buffer and analysed using solution inductively coupled plasma-mass spectrometry (ICP-MS). **B**, Representative microdroplet images illustrating use of laser ablation-ICP-MS (LA-ICP-MS) to measure copper levels in soluble and insoluble fractions of whole homogenates. **C**, Copper in the soluble fraction of whole homogenates of human spinal cord measured using LA-ICP-MS. **D**, Copper in the insoluble fraction of whole homogenates of human spinal cord measured using LA-ICP-MS. Symbols shown in A, C and D represent individual controls and ALS cases. Lines in A, C and D represent mean ±S.E.M. P values represent significance.

**Figure 2.**
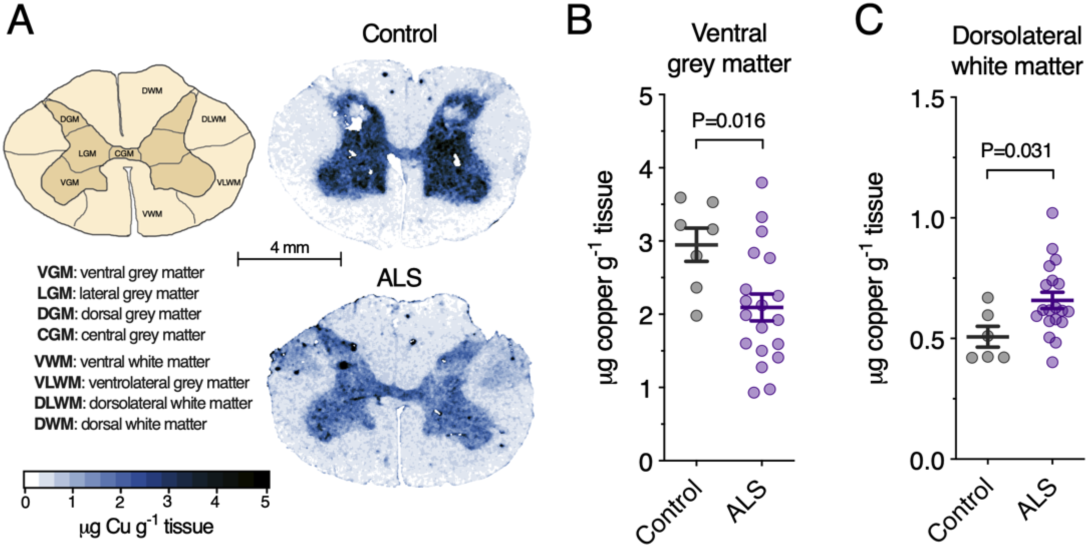
Copper in sporadic ALS-affected spinal cord relative to anatomical distribution. **A,** Schematic illustration of anatomical regions of interest in transverse plane of human spinal cord sections and representative anatomical heatmaps for copper as measured using laser ablation-inductively coupled plasma-mass spectrometry (LA-ICP-MS). **B**, Copper content in ventral grey matter region of interest in human spinal cord measured using LA-ICP-MS. **C**, Copper content in dorsolateral white matter region of interest in human spinal cord measured using LA-ICP-MS. Results for additional regions of interest are shown in Table 2. Symbols shown in B and C represent individual controls and ALS cases. Lines in B and C represent mean ±S.E.M. P values represent significance.

### Altered gene expression in sporadic ALS and mutant SOD1 mice

Physiological distribution of copper within cells involves co-ordinated function of a broad range of copper transporters and chaperones. In light of the prominent copper changes detected in spinal cord grey matter (Fig. 2, Table 2), we measured expression of 20 genes associated with cellular copper handling to gauge the extent to which these copper handling/delivery pathways may be affected in sporadic ALS. These analyses revealed increased expression of 12 of the 20 genes analysed within the ALS-affected spinal cord grey matter (Fig. 3A). Assessment of a homologous panel of genes in spinal cord extracts from SOD1^G37R^ mice revealed increased expression of a similar magnitude for eight out of 16 genes (Fig. 3B). Four of the differentially expressed genes detected in the human tissue (*STEAP4*, *MT1A*, *MT2A*, *XIAP*) were also significantly altered in the mouse model (*Steap4*, *Mt1*, *Mt2*, *Xiap*). Overall quantitation of the molecular signature for copper handling indicated similar changes in spinal cord from human cases of sporadic ALS compared to spinal cord from the mutant SOD1 transgenic mouse model (Fig. 3C,D).

**Figure 3.**
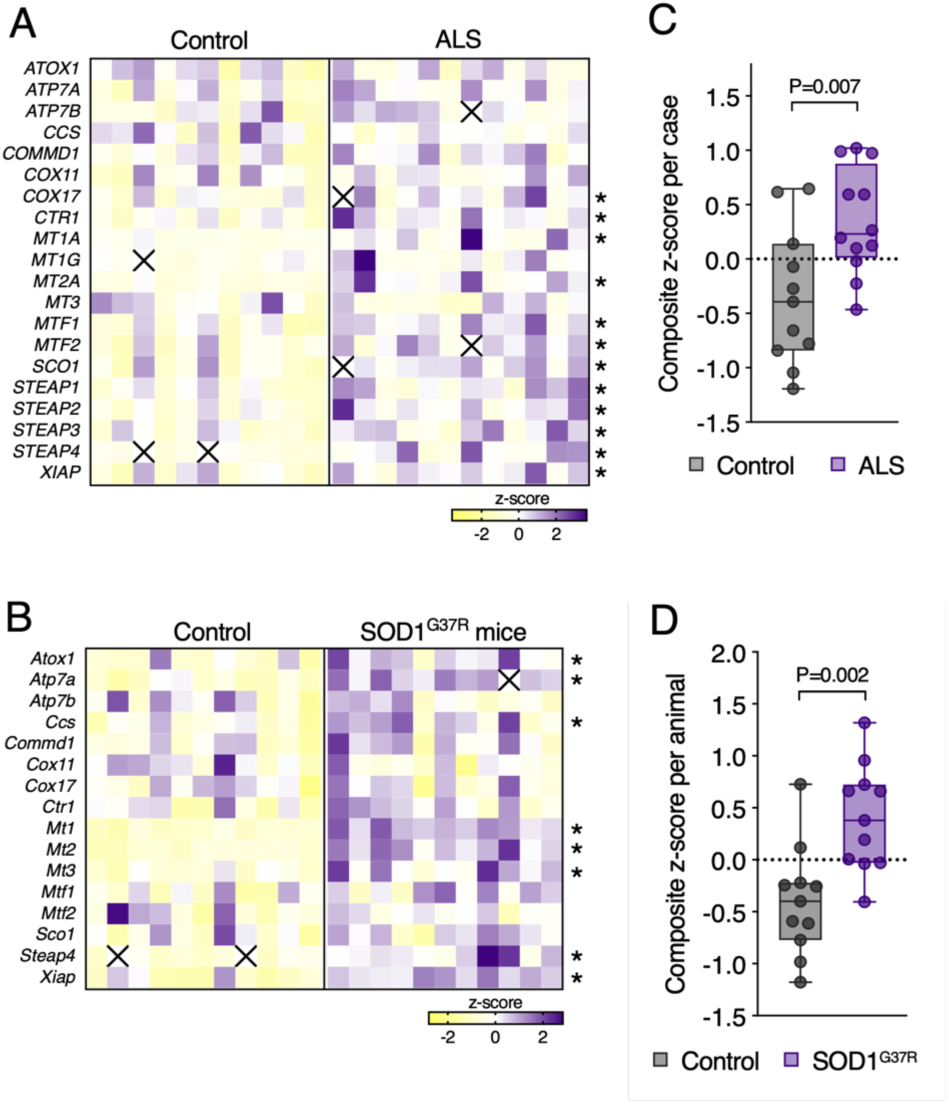
Molecular signature of copper imbalance in sporadic ALS. **A**, Z-score heatmap showing expression of genes encoding copper transporters and chaperones in human grey matter spinal cord measured by quantitative RT-PCR. **B**, Z-score heatmap showing expression of genes encoding copper transporters and chaperones in whole homogenate mouse spinal cord measured by quantitative RT-PCR. ALS model mice are transgenic SOD1G37R mice and controls are non-transgenic littermates. **C**, Composite z-score for overall copper handling in individual control and ALS cases shown in A. **D**, Composite z-score for overall copper handling in individual control and SOD1G37R mice shown in B. Composite z-scores are the average z-score value for each gene measured per individual case or animal (i.e. 20 genes per human case, 16 genes per animal). Squares and symbols in all panels represent individual controls and ALS cases or animals. Box and whisker plots in C and D represent median, 25^th^ and 75^th^ percentiles, and min-max values. P values represent significance and asterisks represent P<0.05.

### Cuproenzyme function is differentially altered in sporadic ALS affected spinal cord

SOD1 and ceruloplasmin both accumulate in copper-deficient states in spinal cords of mutant SOD1 mice^1,3,5,11^. Here, we assessed these cuproenzymes, and additional cuproenzymes, DβH and lysyl oxidase, in human, sporadic ALS-affected spinal cord. Ceruloplasmin and its homolog hephaestin promote iron efflux from certain cells via their multi-copper ferroxidase activity^19^. Here, consistent with mutant SOD1 mouse models^11^, we found that total ferroxidase activity was decreased in ALS (Fig. 4A) even though ceruloplasmin protein levels were increased (Fig. 4B), consistent with an increase in copper-deficient (inactive) ceruloplasmin in sporadic ALS.

**Figure 4.**
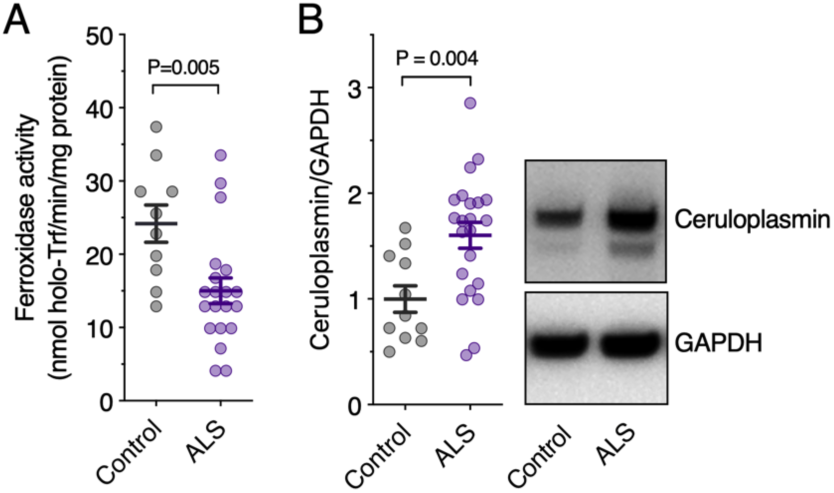
Ferroxidase activity and ceruloplasmin levels in sporadic ALS. **A**, Ferroxidase activity in TBS-insoluble fraction of whole homogenate of human spinal cords measured as the rate of holo-transferrin produced min^-1^ mg^-1^ protein. **B**, Ceruloplasmin protein levels in human spinal cord determined by western blot. Ceruloplasmin protein levels are normalised to the loading control GAPDH. Representative western blot images are shown. Symbols in A and B represent individual controls and ALS cases. P values represent significance.

**Figure 5.**
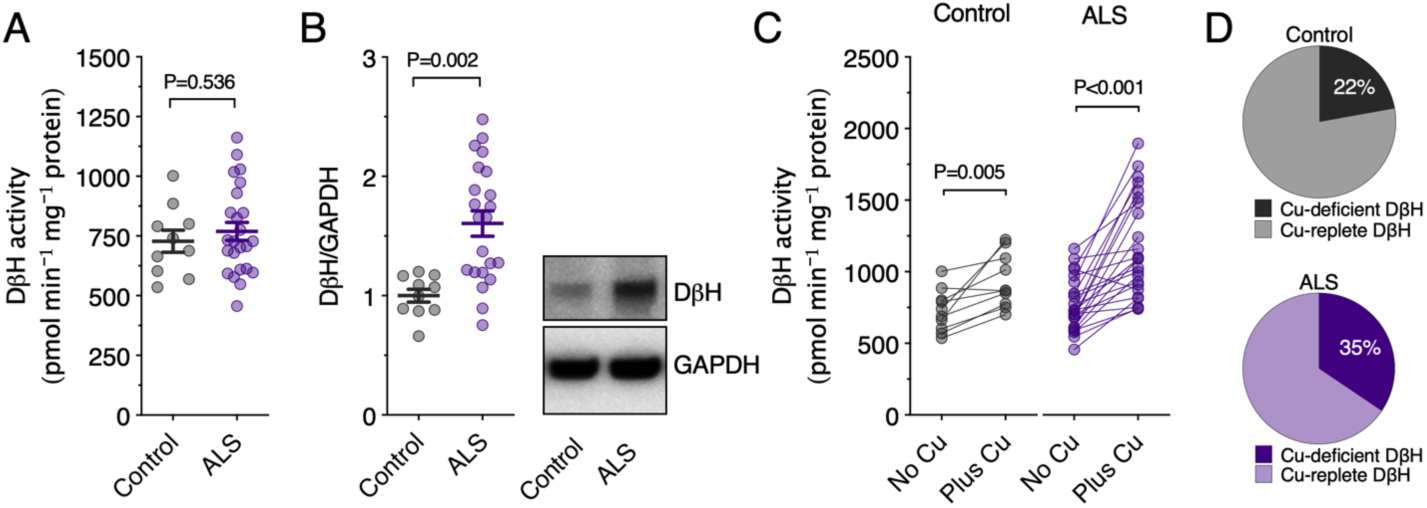
Dopamine β-hydroxylase (DβH) activity and levels in sporadic ALS. **A**, DβH activity in TBS-insoluble fraction of whole homogenate of human spinal cords measured as the rate of norepinephrine produced min^-1^ mg^-1^ protein. **B**, DβH protein levels in human spinal cord determined by western blot. DβH protein levels are normalised to the loading control GAPDH. Representative western blot images are shown. **C**, DβH activity determined as per A with additional reactions for each sample conducted after copper supplementation. **D**, Percentage of copper-deficient and copper-replete DβH in controls and ALS cases derived from data in C. Symbols in A, B and C represent individual controls and ALS cases. P values represent significance.

In contrast to ceruloplasmin, SOD1 protein levels and SOD1 activity were unchanged (SOD1 protein: 1.00±0.04 in control, 0.99±0.05 in ALS, P=0.88; SOD1 activity: 39.42±2.31 % inhibition of pyrogallol oxidation in control, 42.73±1.37 in ALS, P=0.21).

Similar to ceruloplasmin, our assessment of DβH also revealed a disconnect in ALS between abundance of the cuproenzyme and its copper-dependent activity, with higher DβH protein levels not matched by a commensurate increase in activity (Fig. 5A,B). Supplementing the *in vitro* activity assay with exogenous copper increased measurable activity in control cases, but more so in ALS cases (Fig. 5C), indicating that while control cases maintain a pool of copper-deficient DβH, this pool is significantly greater in sporadic ALS (Fig. 5D). In contrast to ceruloplasmin and DβH where copper availability in ALS appears rate-limiting, lysyl oxidase activity was higher in ALS cases compared to controls, despite unchanged protein levels (Fig. 6A-C), indicating greater copper loading onto this cuproenzyme in ALS.

**Figure 6.**
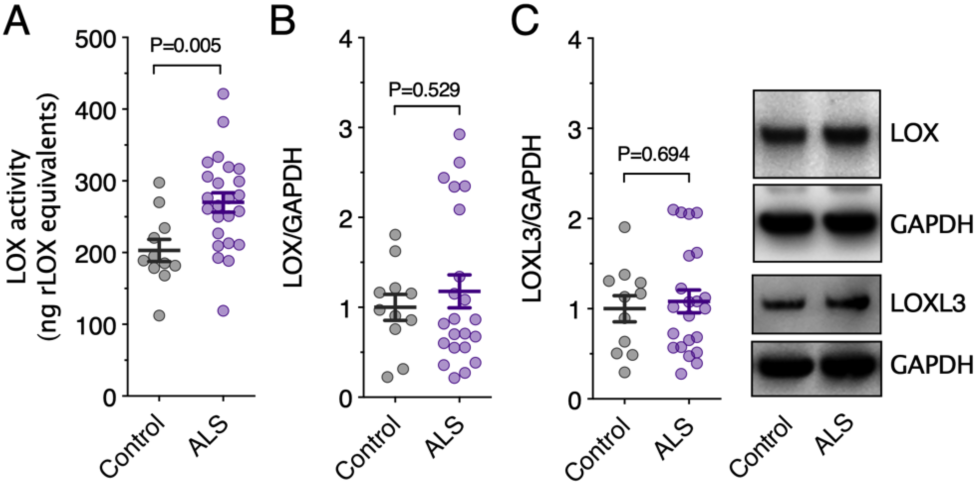
Lysyl oxidase (LOX) activity and protein levels in sporadic ALS. **A**, Lysyl oxidase activity in TBS-insoluble fraction of whole homogenate of human spinal cords measured as the rate of resorufin produced relative to recombinant lysyl oxidase (rLOX) standards. **B**, Lysyl oxidase protein levels in human spinal cord determined by western blot. **C**, Lysyl oxidase homologue 3 (LOXL3) protein levels in human spinal cord determined by western blot. Lysyl oxidase and lysyl oxidase homologue 3 protein levels are normalised to the loading control GAPDH. Representative western blot images are shown. Symbols in A, B and C represent individual controls and ALS cases. P values represent significance.

## DISCUSSION

Transgenic mice expressing mutant SOD1 are the most robust animal model available for ALS. However, no therapeutic agent developed and tested in these animals has successfully translated to disease-modifying therapy in patients. While this has led some to question the utility of mutant SOD1 models, it has also prompted interrogation of the reproducibility of reported pre-clinical outcomes. To this end, the ALS Therapy Development Institute has independently assessed prior reports of pre-clinical efficacy in mutant SOD1 mice. To date, the copper-containing compound Cu^II^(atsm) remains as the first and only drug candidate that the ALS Therapy Development Institute has been able to validate^6^. This provides strong support for Cu^II^(atsm), but validation of pre-clinical studies involving mutant SOD1 models does not address a remaining significant challenge in ALS drug development, namely, that only ~2% of ALS cases in the clinic are caused by SOD1 mutations. The majority of cases are sporadic with an unknown aetiology. The translational relevance of therapeutic outcomes from mutant SOD1 mice to the treatment of sporadic ALS is of concern.

Here, we show that improving copper availability to cuproenzymes in the CNS may represent a therapeutic mechanism of action that links pre-clinical outcomes in mutant SOD1 mice to sporadic ALS. Direct evidence for disrupted copper availability in sporadic ALS is provided by our *in situ* quantitation of copper using LA-ICP-MS (Fig. 2, Table 2). Most conspicuously, these analyses revealed ventral grey matter, the primary anatomical site of pathology in ALS, as the spinal cord region with the greatest decrease in copper.

Consistent with these changes impacting the physiological requirement for copper, we show altered expression for the majority of genes involved in copper handling (Fig. 3A), resulting in an overall molecular signature for copper handling that is shifted in sporadic ALS (Fig. 3C). Significantly, we show that mutant SOD1 mice model this molecular feature of the sporadic disease in humans (Fig. 3B,D). Evidences exist for these transcriptional changes manifesting in altered abundance of the associated proteins^20,21^. Moreover, previous reports have demonstrated that cuproenzymes in the spinal cord of mutant SOD1 mice do not obtain their requisite supply of copper, and that genetic or pharmacotherapeutic release from this bottleneck improves the animals’ ALS-like phenotype^1,3,5,11^. The most explicit evidence for this came from a study in which mutant SOD1 mice were treated with Cu^II^(atsm) enriched with the natural copper isotope ^65^Cu. Administration of ^65^Cu^II^(atsm) followed by liquid chromatography-ICP-MS analysis confirmed that copper from the orally administered Cu^II^(atsm) entered the bioavailable pool of copper in the CNS, resulting in subsequent incorporation into endogenous cuproproteins^3^, resulting in improved activity of the cuproenzymes that accumulate in a copper-deficient state in mutant SOD1 mice^1,3,5,11^.

Here, we provide evidences for the cuproenzymes ceruloplasmin and DβH both accumulating in copper-deficient states in sporadic ALS (Fig. 4 and 5). Although iron can be exported through other mechanisms, efflux via ferroportin, which requires oxidation of iron by ceruloplasmin, constitutes a major pathway in some cells. Thus, decreased copper-dependent ferroxidase activity could contribute to the accumulation iron, which has been identified in ALS patients by magnetic resonance imaging^22^. Similarly, catecholamines are essential for neuronal function and DβH is an important part of their regulation because it converts dopamine to norepinephrine. The data we present here provide evidence for potential disruption in catecholamine levels in ALS being related to altered copper bioavailability. An unsatiated requirement for copper in these cuproenzymes may therefore contribute to ALS. Delivery of copper from Cu^II^(atsm) to cuproenzymes unsatiated for their natural copper requirement has been demonstrated in mutant SOD1 mice^3,5^. In these animals, the treatment also protects motor neurones in the CNS, mitigates associated deterioration of motor function, and improves overall survival^1–7^.

Notably, our analyses indicate copper perturbations in sporadic ALS cannot be broadly characterised as an overall copper insufficiency, indicating that ALS is not caused by simple tissue nutritional copper deficiency. This is apparent in our assessment of copper partitioning into TBS-soluble and -insoluble fractions of the spinal cord, with loss of copper from the soluble fraction and a concomitant increase in the insoluble fraction (Fig. 1C,D). Additionally, although ceruloplasmin and DβH are proportionately copper-deficient, copper-dependent lysyl oxidase activity is increased in sporadic ALS (Fig. 6A). Lysyl oxidase and its homologues catalyse cross-linking of extracellular matrix proteins^23^. Aberrant changes to the extracellular matrix have been described in ALS that may adversely affect CNS structure and integrity of the blood brain barrier^24^. Increased LOX activity could contribute to these changes through its effect on important constituents of the extracellular matrix such as collagen IV. Overall, it is possible that copper perturbations in ALS involve an internal redistribution that might benefit some cuproenzymes at the expense of others.

Cu^II^(atsm) labelled with radioactive isotopes of copper is utilised as a positron emission tomography (PET) imaging agent, with prominent use in imaging hypoxic tumour^25,26^. Other studies have explored its application to imaging diseases of the CNS^27,28^, including sporadic ALS where motor cortical accumulation of the PET signal correlates with symptomatic stage of disease progression determined using the ALS Functional Rating Scale (Revised)^29^. The specific factors that drive the observed localised accumulation in ALS are not yet confirmed. However, redox imbalance involving a hyper-reductive state is implicated^30,31^. In brief, cellular retention of copper from bis(thiosemicarbazonato)-copper^II^ compounds such as Cu^II^(atsm) is dictated, in part, by intracellular reduction of the copper followed by its dissociation from the ligand. Electron donating methyl groups on the atsm ligand confer a reduction potential to Cu^II^(atsm) which means the copper is relatively resistant to intracellular reduction under physiological conditions. However, under hypoxic conditions electron flux through the mitochondrial electron transport chain is impeded, resulting in a hyper-reductive state that promotes reduction of the copper in Cu^II^(atsm) and its dissociation from the complex. Significantly, impaired electron flux through the electron transport chain even under normoxic conditions is sufficient to generate the hyper-reductive state that promotes reduction and dissociation of Cu^II^(atsm) within cells^30,31^. Mitochondrial dysfunction and/or tissue hypoxia in ALS^32^ may provide conditions within the disease-affected tissue that promote selective release of copper from Cu^II^(atsm). The presence of an aberrant pool of copper-deficient cuproenzymes within the same tissue will also promote localised accumulation of the copper through binding to cuproproteins which is a determinant of cellular retention of the copper^33^. The preferential release and retention of copper from Cu^II^(atsm) provides a plausible explanation for a PET signal in ALS patients that highlights the affected motor cortex^29^.

Additional mechanisms for the neuroprotective activity of Cu^II^(atsm) have been described. The compound exhibits potent anti-inflammatory effects^34,35^, it inhibits ferroptosis, an iron-dependent form of non-apoptotic cell death^36^, and it protects against peroxynitrite-driven toxicity^37^. All of these protective activities are pertinent to aberrant processes associated with ALS and therefore may also contribute to the potential therapeutic mechanism of Cu^II^(atsm) in treating ALS. Moreover, these potential therapeutic targets are also pertinent to other neurodegenerative diseases. Cu^II^(atsm) treatment results in neuroprotection and mitigation of symptoms of neuronal decline in four different animal models of Parkinson’s disease (PD)^37^, leading to phase 1 clinical assessment of the compound in PD patients^38^. And as in ALS patients, monitoring radio-labelled Cu^II^(atsm) in PD patients via PET imaging shows selective accumulation in the disease-affected basal ganglia^28^. As above, this selective accumulation implies region-specific influences of a hyper-reductive state in PD, a possibility consistent with evidences for mitochondrial dysfunction in PD^39^. It is also consistent with evidences for copper insufficiency in the PD-affected brain, including decreased copper levels, decreased ceruloplasmin ferroxidase activity, and the accumulation of copper-deficient SOD1^40–42^.

Cu^II^(atsm) has produced favourable outcomes from an open-label phase 1 trial in which the ALS Functional Rating Scale (Revised), Edinburgh Cognitive and Behavioural ALS Screen, and seated forced vital capacity were included as secondary outcome measures^43,44^. A blinded, placebo-controlled phase 2/3 trial is now underway^45^. Pre-clinical support for Cu^II^(atsm) in the treatment of ALS is primarily derived from *in vivo* studies involving mutant SOD1 animal models of the disease^1–7^, raising questions concerning applicability to the preponderant amount of cases which does not involve mutant SOD1. Here, we show that disrupted copper availability is a feature of sporadic ALS, recapitulating the unsatiated copper requirement of the mutant SOD1 mice. Therefore, the benefits of Cu^II^(atsm) treatment and imaging can be expected in sporadic cases of ALS, and not just those involving mutant SOD1.

## ACKNOWLEDGEMENTS

Human tissue samples were obtained from the Victorian Brain Bank (Florey Institute of Neuroscience and Mental Health, the University of Melbourne, Australia) with assistance from Ms Fairlie Hinton and Mr Geoff Pavey, and also the MS Society Tissue Bank (Wolfson Neuroscience Laboratories, Imperial College London, United Kingdom) with assistance from Dr Djordie Gveric. Total copper levels in human spinal cord were measured at the Biometals Facility at the Florey Institute of Neuroscience and Mental Health, the University of Melbourne, by Ms Irene Volitakis. This research was supported by funding from the Motor Neurone Disease Research Institute of Australia (Beryl Bayley Fellowship to JBWH; Betty Laidlaw MND Research Project to PJC, BRR, DJH, ARW, JSB, PSD and CAM; Jenny Barr Smith MND Research Project to PJC, AIB, BRR, JSB and CAM) and the University of Melbourne.

## AUTHOR CONTRIBUTIONS

JBWH, and PJC contributed to conception and design of the study, acquisition and analysis of data, and drafting the manuscript. JRL and AIB contributed to design of the study, acquisition and analysis of data, and drafting the manuscript. CR, SWM, BP, CAM, JSB, ARW, PSD, DJH and BRR, contributed to acquisition and analysis of data.

